# Long-term Antibody Release Polycaprolactone (PCL) Capsule and the Release Kinetics In Natural and Accelerated Degradation

**DOI:** 10.1101/2022.06.06.493286

**Authors:** Waterkotte Thomas, Xingyu He, Apipa Wanasathop, Kevin Li, Yoonjee Park

## Abstract

Although therapy using monoclonal antibodies (mAbs) has been steadily successful over the last 20 years, the means of delivery of mAbs has not been optimized, especially for long-term delivery. Frequent injections or infusions have been current standard of care. In this study, we have developed a long-term antibody biodegradable implant using a porous polycaprolactone (PCL) capsule. It released Bevacizumab (Bev) slowly for 8 months to date. The Bev release kinetics fit a drug release model with experimental data of the diffusion coefficient and partition coefficient through the polymer capsule. Since screening drug release profiles for the long-term (> 6 months) is time consuming, an accelerated degradation method was used after validating characteristics of the PCL capsule in natural and accelerated degradation conditions. The correlation of time period between the natural and the accelerated degradation was determined. Overall, the study suggests mAbs can be released from a porous PCL capsule without an effect of the polymer degradation over the long period (~ 6 months) and the long-term release kinetics can be determined by the accelerated degradation within 14 days.

## 1. Introduction

Monoclonal antibodies (mAbs) are the largest class of biopharmaceuticals and are used in the treatment of many chronic diseases including cancer, immunological disorders, and infectious diseases.^1^ US Food and Drug Administration (FDA) and European Medicines Agency (EMA) approved 36 mAb drugs from 2013 to 2017, and the FDA approved 23 new mAbs in 2018-2020.04. ^2^ The size of the global mAbs market was worth USD 39.1 billion in 2021 and this figure is estimated to be growing and will reach USD 50.62 billion by 2026. This is because the number of patients treated by mAbs is increasing as well as the possibility of more mAbs coming to market is rising. ^3^

Therapeutic mAbs have been widely developed and used for chronic disease treatment. For example, bevacizumab or ranibizumab have been used for chronic wet age-related macular degeneration (AMD) or various types of cancer. Treatment of the diseases involves the suppression of the vascular endothelial growth factor (VEGF) protein. Current standard of care for AMD is repetitive monthly intravitreal injections of these mAbs directly to the eye. The frequent injections are necessary partly because of a relatively short half-life of the drugs in the target sites. Bevacizumab (Bev) administered intravitreally has a half-life of 5 to 10 days. ^4^ However, the frequent intravitreal injections can cause issues, including hemorrhage, corneal erosion, and post-injection ocular surface discomfort (i.e. burning, stinging, or itching). ^5^ In addition, this is a significant burden to the patient, the patient’s family, and the healthcare system.

There is no biodegradable drug delivery system for the long-term sustained antibody release, i.e. over a month, to date. Although a port delivery device has been recently launched for delivery of ranibizumab for posterior eye disease treatment, long-term durability and whether it should be replaced after a certain duration of time as well as the appropriate procedure for safe removal of the implant should complications arise are uncertain. ^6^ The port delivery system is surgically inserted via the pars plana area in the superotemporal quadrant of the eye and releases antibodies sustainably for several months. In addition, the port delivery system has been associated with a 3-fold higher rate of endophthalmitis than monthly intravitreal injections.

There are biodegradable implants available for small drug molecules, such as steroid or hormone designed for the long term, i.e. 6 months. However, it has been challenging to develop a system that releases antibodies with a molecular weight of ~150 kDa, which are much bigger than typical drug molecules with several hundred Da. The typical method of implant fabrication is a melt-extrusion method, which requires high temperature (100~200 °C) to melt polymers. ^7^ Although some drug molecules are stable at the high temperature and can be mixed with the polymer to be a matrix-type implant, antibodies are not stable at the high temperature and this fabrication method cannot be used for antibody implant fabrication.

Here, we have developed a nanoporous capsule implant that releases antibodies slowly over 200 days, which are degradable in the body utilizing polycaprolactone (PCL). PCL is an FDA approved biodegradable polymer for use in implantable biomaterials and injectable implants. ^8^ The slow degradation and hydrophobicity of PCL is one reason why it is extensively used in long-term drug delivery systems. ^9^ In this study, PCL will be used to develop Bev (antibody) implants for a long-term biodegradable drug delivery platform.

In addition, degradable polymers are known to undergo morphological changes over degradation and these changes may affect the antibody release kinetics from the implant. For examples, pore size in the polymer increases over degradation and surface erosion can reduce the thickness of polymer, which may impact on faster antibody release from the implant. Thus, this study will determine the antibody release kinetics for the long-term and whether the kinetics is affected by the degradation. Because degradation rate of PCL is slow, which takes at least 6 months and up to 4 years for noticeable degradation, the release kinetics and the degradation of the implant will be determined in an accelerated condition in this study. The polymer degradation proceeds via mainly three pathways, i.e. biological, chemical or physical means. In vivo and in vitro degradation were reported to occur at the same rate, suggesting no significant contribution by enzymes initially.^10 11^ Biological degradation was more pronounced when M_n_ (molecular weight) reaches ~5000 for PCL.^12^ The main mode of degradation for high molecular-weight aliphatic polyesters is hydrolytic random scission. ^13^ Thus, accelerated degradation using an acidic or basic medium, which enhances the hydrolysis of polyester, mimics physiological conditions better than other methods, such as temperature acceleration. A basic medium, such as a sodium hydroxide (NaOH) solution, has been widely studied for accelerated polymer hydrolysis/degradation.^9 14^ Thus, the accelerated degradation method will be useful to determine the drug release kinetics over degradation. Several publications including ASTM International (American Society for Testing and Materials) still suggest comparing the morphological structure between natural and accelerated degradation.

In this article, we will present nanoporous PCL capsule implants developed for long-term antibody release as shown in the schematics of **Figure 1** with three different pore sizes and drug release rates in the accelerated degradation conditions for the first time. Furthermore, the antibody release kinetics will be fit to mathematical models using permeability and partition coefficient values determined by experiment. In addition, this study will evaluate the effectiveness of the accelerated system for predicting the long-term drug release kinetics with characterization of the morphologies of the PCL capsule both in natural and accelerated degradation conditions.

**Figure 1.**
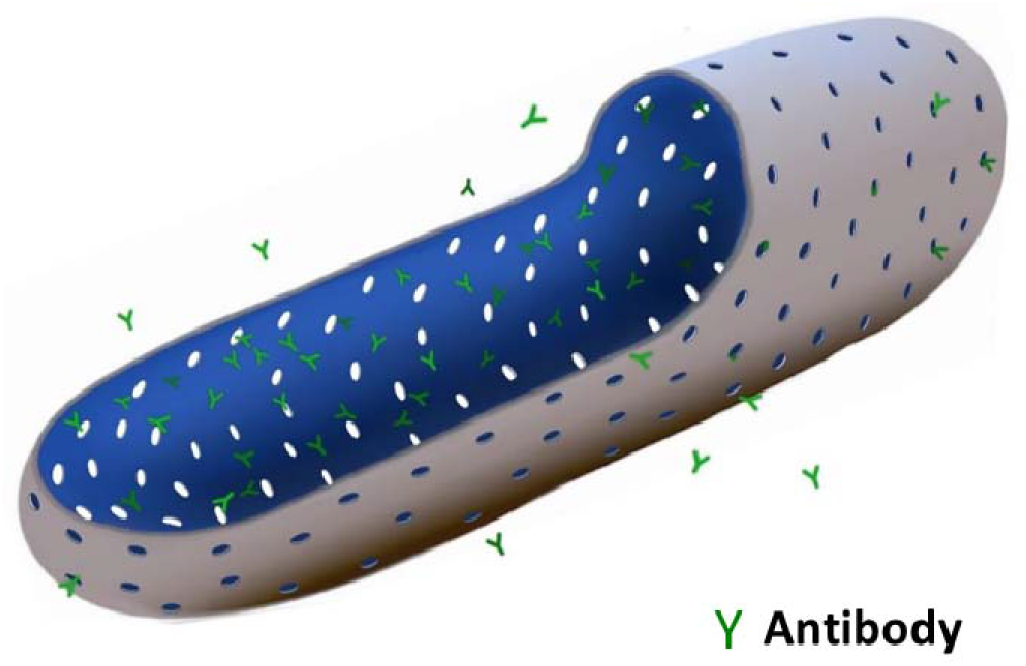
Schematics of the nanoporous PCL antibody implant.

## 2. Materials and Methods

### 2.1. Materials

Poly(caprolactone), PCL, (MW 65,000-75,000) was purchased from PolyScitech, Inc. (West Lafayette, IN). Dichloromethane (DCM), sodium hydroxide (NaOH), and polyethylene glycol (PEG, Average MW 3350) were purchased from Fisher Chemical (USA). Bevacizumab (Bev) solution was purchased from Pfizer (Zirabev) (New York, NY).

### 2.2. PCL sheet synthesis

The nano porous PCL sheet was synthesized by first creating a 50 mg/mL PCL solution in DCM with varying amounts of PEG as a porogen for differing porosities. The ratios were 0.0, 0.05 and 0.1 PEG to PCL by weight. 800 μL of the PCL solution was placed in a mold floating in a bath sonicator at 15°C. The mold was covered with parafilm and sonicated at 50% power for 80 minutes to create a homogeneous distribution of porogen and slowly evaporate the DCM. After sonication, the sheet was removed, and air dried in a chemical hood at room temperature overnight.

### 2.3. Implant fabrication

To create PCL implants, the PCL sheet was cut into 0.5 x 1 cm^2^ pieces using a razor blade. The pieces were then rolled around a 22-gauge needle at 45°C to create a double layered tube 1 cm in length and 0.0946 cm in diameter. The tube was then submerged in deionized water overnight to leach out the PEG and create a porous structure and then dried overnight at room temperature. Once dried, a 60°C iron was used to clamp one end of the tube shut and a 26-gauge needle was used to load 4 μL of a Bev solution into the tube. Once the Bev solution was loaded, the iron was used to seal the other end shut and then both ends were superglued to prevent any leakage that may occur. The whole implant fabrication process is shown in **Figure 2.**

**Figure 2.**
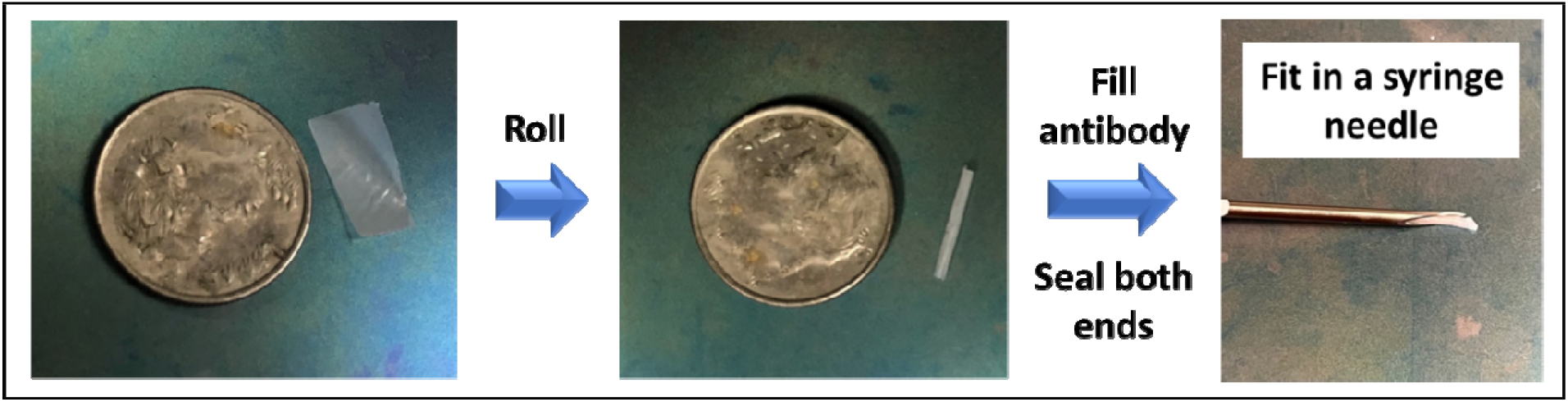
Implant fabrication process (20-gauge needle pictured).

### 2.4. Drug release profile

Implants loaded with the Bev solution were placed in 1 mL of PBS and incubated at 35°C to mimic the temperature of the human eye (n = 3). The PBS solution was removed and replaced with fresh PBS every 24 h, 72 h, or 7 days depending on how long the implant has been incubated. The removed solution was analyzed using UV-Vis at 280 nm to determine the Bev concentration. Using the concentration, the amount of Bev released was plotted to determine the release profile.

### 2.5. Accelerated drug release profile

The implants used in these experiments were prepared in the same way as section 2.4. Instead of placing the implants in 1 mL of PBS, the implants were placed in 1 mL of 0.1 M NaOH at 35°C for accelerated release (n = 3). Every 24 h the NaOH solution was removed, analyzed using UV-Vis at 285 nm due to the lambda max shift in Bev in NaOH, and fresh NaOH was added. The study was conducted over 14 days due to the breakdown of 0.1 PEG PCL implants. The concentration found was used to determine the amount of Bev released each day and then plotted.

### 2.6. SEM Imaging

The surface and cross-section structures were analyzed using scanning electron microscopy (SEM) (Thermo Fisher Scios Dual Beam SEM, Hillsboro, OR). The PCL sheet or tube was placed on a SEM sample holder with dual sided graphite conductive tape. The samples were sputter-coated with 5 nm platinum/gold nanoparticles for 10 s using Denton Vacuum Desk II (Moorestown, NJ). To obtain the cross-section view, tubes were submerged in glycerol, placed in liquid nitrogen until frozen, and then fractured to get the best cross-section view without having the edges roll from cutting at room temperature. The samples were then rinsed with DI water and dried overnight. The mean pore sizes and pore density were analyzed using ImageJ (NIH).

Porosity was calculated using the following equation:

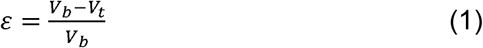

where *V_b_* is the bulk volume (volume of membrane if no pores) and *V_t_* is the true volume of the membrane (volume of membrane taking out pore volume). The bulk volume, *V_b_*, was calculated using the thickness of the membrane determined by SEM and the area of the polymer sheet used to roll the implants. The true volume, *V_t_*, is the bulk volume, *V_b_*, minus the total volume of the pores in the membrane. The volume of pores was determined by finding the volume of a pore using the average pore diameter (determined by SEM) and assuming spherical pores, the pore density (pores/nm^3^ determined by SEM), and the total volume of the implant. Multiplying the pore density and volume gives the total number of pores which multiplied by the average pore volume give the total volume of pores.

### 2.7. Permeability

The permeability experiments were conducted using a vertical Franz Diffusion cell. The donor side was filled with 200 μL of 50 mg/mL Bev solution and the receiver side was filled with 5 mL of PBS. The membrane used was a 1cm x 1cm double layered PCL sheet. The membrane was made by placing two 1 x 1 cm PCL sheets on top of each other and pressing them together at 45°C to create a double layered membrane. The membranes were soaked in DI water for 24 h to remove the PEG and then placed in the cell without drying. Once the membrane was securely placed between the donor and receiver chambers 200 μL of the donor solution was loaded and then the opening was sealed with parafilm to reduce evaporation altering the concentration of the donor and receiver solutions. 500 μL of the receiver and 10 μL of the donor solution were removed every 72 h to determine the concentration using UV-Vis. The removed solutions were replaced by fresh PBS for the receiver solution and the donor was replaced by fresh donor solution or completely replaced depending on the remaining concentration.

After 12 days, the mass of Bev that permeated the membrane was plotted against time to get *dQ/dt* in the equation:

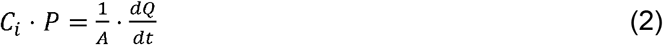

where *C_i_* is the initial concentration (donor solution concentration), *P* is the permeability coefficient, *Q* is the amount of Bev permeated at time *t,* and *A* is the area of the exposed membrane.

### 2.8. Partition Coefficient

The partition coefficient *K* is required for the release kinetics model equation. To measure the partition coefficient, a 0.5 x 0.5 cm^2^ double-layered PCL membrane was made in a similar way to the membranes in the permeability experiment. The membranes were soaked in 200 μL of PBS for 48 h. After 48 h the membranes were blotted with a Kimwipe to remove surface liquid and then weighed to determine the wet mass of the membrane. The membranes were then dried overnight in a chemical hood. The dry membranes were then placed in 200 μL of an equilibrium Bev solution for 72 h to allow the membranes to absorb the Bev from the solution. After 72 h the membranes were removed from the equilibrium, blotted, and placed in 200 μL of fresh PBS to extract Bev for 48 h. This extraction process was repeated until the amount extracted was 10% of the first extraction. The remaining equilibrium solution and all extraction solutions were analyzed using UV-Vis to determine the Bev concentration. The partition coefficient was then determined using the equation below.

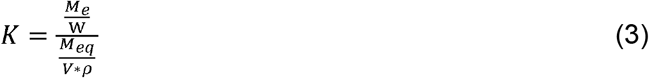

where *M_e_* is the total mass extracted, *W* is the wet mass of the membrane, *M*_eq_ is the amount of Bev left in the equilibrium solution, *V* is the volume of equilibrium solution left, and *ρ* is the density of the solution.

### 2.9. Modealing Equations

The equation used to model the release profile is

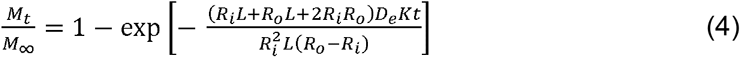

where *M_t_* is the mass of Bev released at time *t, M*_∞_ is the mass of Bev loaded in the implant, *R_i_*, is the inner radius of the implant, *R_i_* is outer radius of the implant, *D_e_* is the effective diffusivity, *K* is the partition coefficient, and *L* is the length of the implant.^15^ This equation models Fick’s law of diffusion of molecules from the lumen of a cylindrical implant through a membrane wall. The permeability of Bev across the PCL membrane can be described by

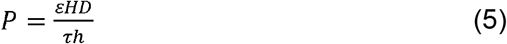

where *ε* is the porosity of the membrane, *H* is the overall hinderance factor for diffusion, *τ* is tortuosity, *h* is the thickness of the membrane, and *D* is the Stokes-Einstein diffusion coefficient:

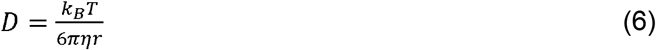

Where *k_B_* is the Boltzmann constant, *T* is the temperature in Kelvin, *η* is the viscosity of water at 25°C, and *r* is the hydrodynamic radius of Bev. ^16^ The overall hindrance factor for diffusion, *H,* is expressed as:

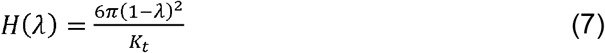

Where λ is the ratio of permeant radius to pore radius and *K_t_*, the hinderance factor, is calculated by:

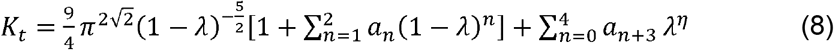

The last value that needs to be determined for Equation 4 is effective diffusivity (*De*). *De* can be determined from the equation:

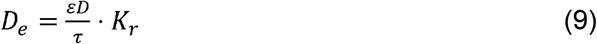

where *K_r_* is the restrictive factor when the ratio of the molecule diameter (*d_m_*) and pore diameter (*d_p_*) is less than 1. The relationship between the diameters is expressed by:

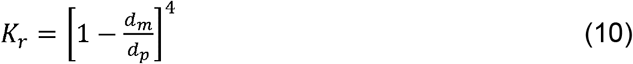

Combining Eq. 5 and Eq. 9 reduces to Eq. 11, which does not require a value for tortuosity.

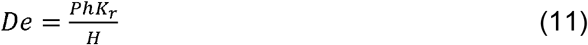

## 3. Results

### 3.1. Characterization of the PCL capsule

PCL sheets with 3 PEG ratios (0.0, 0.05, and 0.1) were synthesized. The overall structure of the implant was imaged using SEM (**Figure 3A**). The scale-like structure on the surface is most likely due to the evaporation of DCM during the sheet preparation. We confirmed that the air side (top) exhibited a scale-like structure (**Figure 3B)** and the mold side (bottom) was smooth except for small indents created by the presence of PEG (**Figure 3C**). The area of scale structure (μm^2^) was significantly different for all 3 PEG conditions. 0.0 PEG, 0.5 PEG, and 0.1 PEG had an average area of 200 ± 10 μm^2^, 5,480 ± 500 μm^2^, and 3,130 ± 220 μm^2^, respectively, indicating PEG affects the surface morphology (p-value <0.05). Along with the scale structure, the top side of the sheet has a high density of pores homogenously distributed across the entire surface (inset of **Figure 3B**). The pore size, porosity, and thickness of the PCL implants with 3 PEG ratios were analyzed using the cross-section SEM images (**Figures 3D–3F, Table 1**). 0.0 PEG PCL sheets exhibited no pores on the surface or in the cross section. The 0.05 PEG ratio had a mean pore size of 520 ± 20 nm, and a porosity of 3.14%. The 0.1 PEG ratio had an average pore size of 620 ± 10 nm and a porosity of 6.88%. The pore size (p = 0.0068) and the porosity (p = 4.8E-4) statistically significantly increased from 0.05 to 0.1 PEG ratio. The average thickness of 0.0 PEG, 0.05 PEG, and 0.1 PEG was 20.5 ± 0.25 μm, 27.6 ± 0.54 μm, and 25.6 ± 0.18 μm, respectively, showing no correlation between the thickness and the PEG ratio.

**Figure 3.**
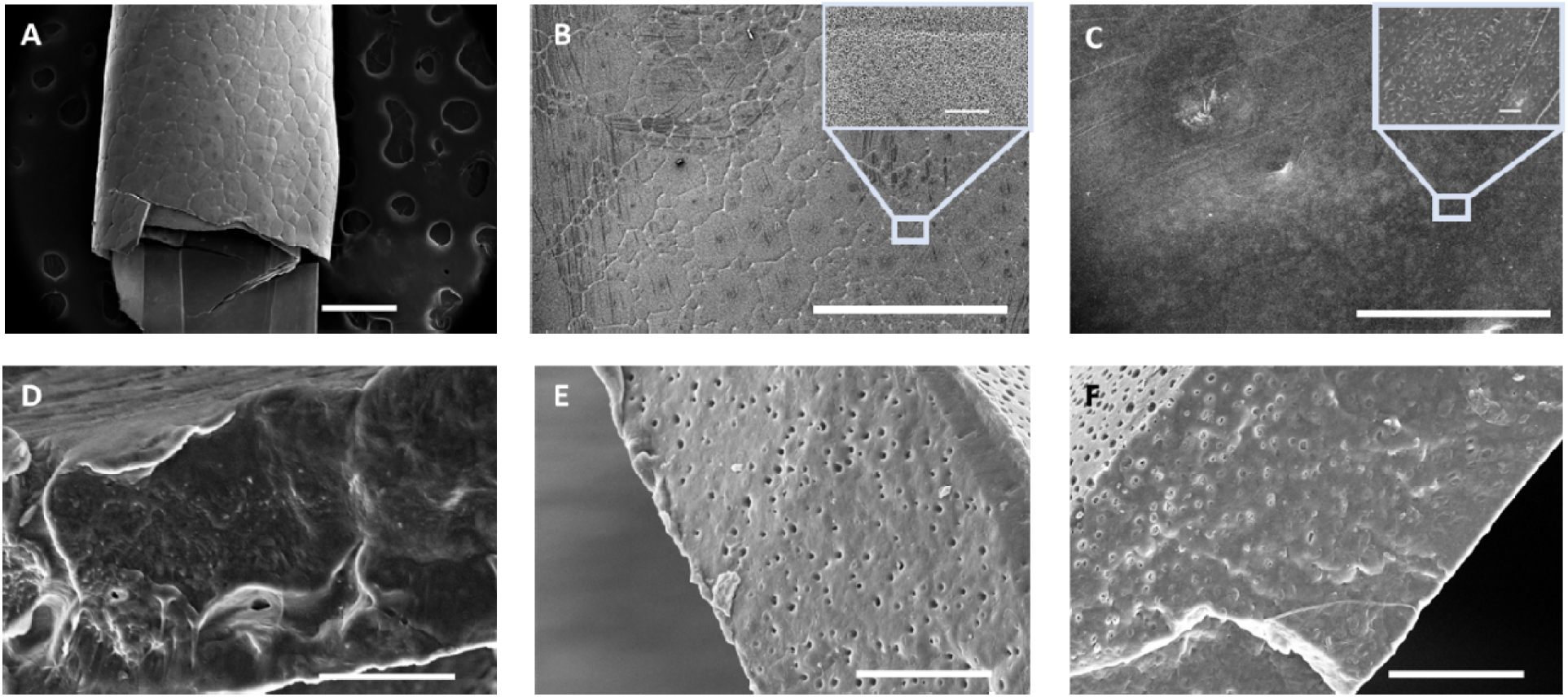
SEM images of (A) 0.05 PEG PCL overall (scale bar 500 μm), (B) 0.05 PEG PCL sheet top, (C) 0.05 PEG PCL sheet bottom, (D, E, F) Cross-sections of 0.0 PEG PCL, 0.05 PEG PCL, and 0.1 PEG PCL, respectively. Scale bars = 10 μm except the overall (A).

**Table 1.** pore size, porosity, scale area and thickness analysis of PCL capsules at Day 0 from SEM cross-section images.

| Day |  | 0.0 PEG | 0.05 PEG | 0.1 PEG |
| --- | --- | --- | --- | --- |
| 0 | Pore Diameter (nm) | N/A | $510 \pm 30$ | $620 \pm 20$ |
| | Porosity (%) | N/A | $3.14 \pm 0.177$ | $6.88 \pm 0.336$ |
| | Scale Area ($\mu\text{m}^2$) | $200 \pm 10$ | $5,480 \pm 990$ | $3,130 \pm 220$ |
| | Sheet thickness ($\mu\text{m}$) | $20.5 \pm 0.25$ | $25.6 \pm 0.18$ | $27.6 \pm 0.54$ |

### 3.2. Drug (Bev) release profile from the PCL implant

The release profile from the implant shows that 97.7% of the Bev was released after 6 months (**Figure 4**). The release was fast in the first ~50 days releasing 75% before slowing down to release 23% in the next 160 days. The decrease in release rate with time is due to the drop in concentration gradient across the membrane as Bev diffuses out of the implant. The release curve shown in **Figure 4** exhibits first-order kinetics which will be discussed further in section 3.6.2.

**Figure 4.**
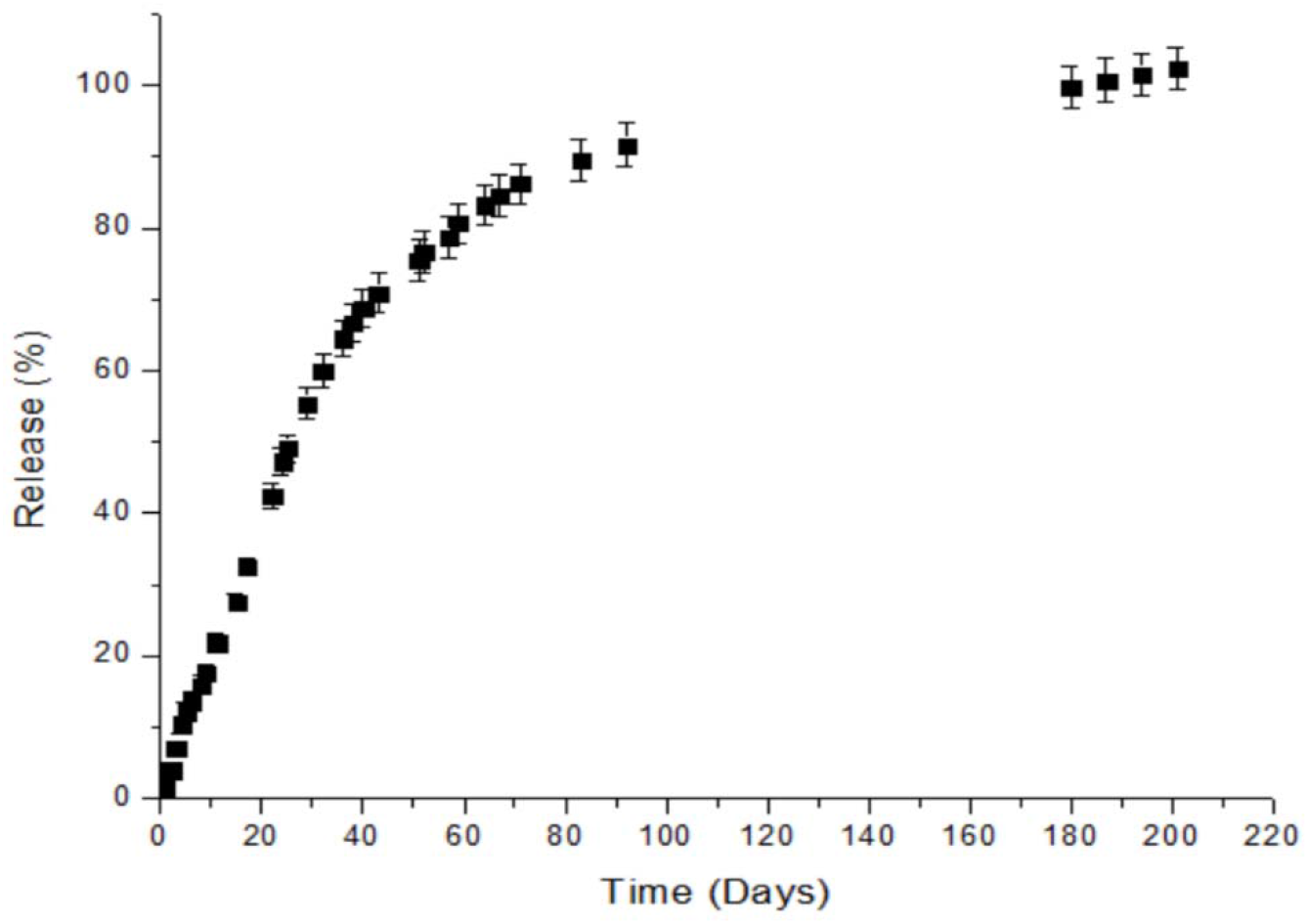
Bev Release in natural condition from 0.05 PEG PCL implants

### 3.3. Drug release profile under accelerated degradation conditions

The Bev release curves from PCL capsules with various PEG ratios were achieved in the accelerated degradation condition utilizing 0.1 M NaOH. The **Figure 5** shows that the time frame of drug release until it showed a plateau reduced to ~2 weeks in the accelerated condition from 6 months in the natural condition. 0.1 M NaOH was chosen to balance between the speed of the drug release over degradation and in-depth analysis of the PCL degradation. Higher concentrations of NaOH (0.25 and 0.5 M) broke down the implants within 2 to 3 days and a lower concentration at 0.05 M had not achieved enough release or degradation within 3 weeks (data not shown). The curves from the 0.05 PEG PCL accelerated release mimic the first-order kinetics from the natural condition release. 0.1 PEG has a 53.4% ± 6.78% burst release in the first 24 hours before slowing down. The 0.0 PEG ratio has a 2-step release curve with a small amount (31.0% ± 7.37%) released in the first 5 days before plateauing until day 10 where the release speeds back up. This second surge of release is due to micro cracks forming in the implant due to degradation, which might have happened earlier for 0.05 and 0.1 PEG, as discussed in section 3.5. Using the curve shapes, a time correlation can be obtained to create a relationship between time in 0.1 M NaOH and time in natural condition.

**Figure 5.**
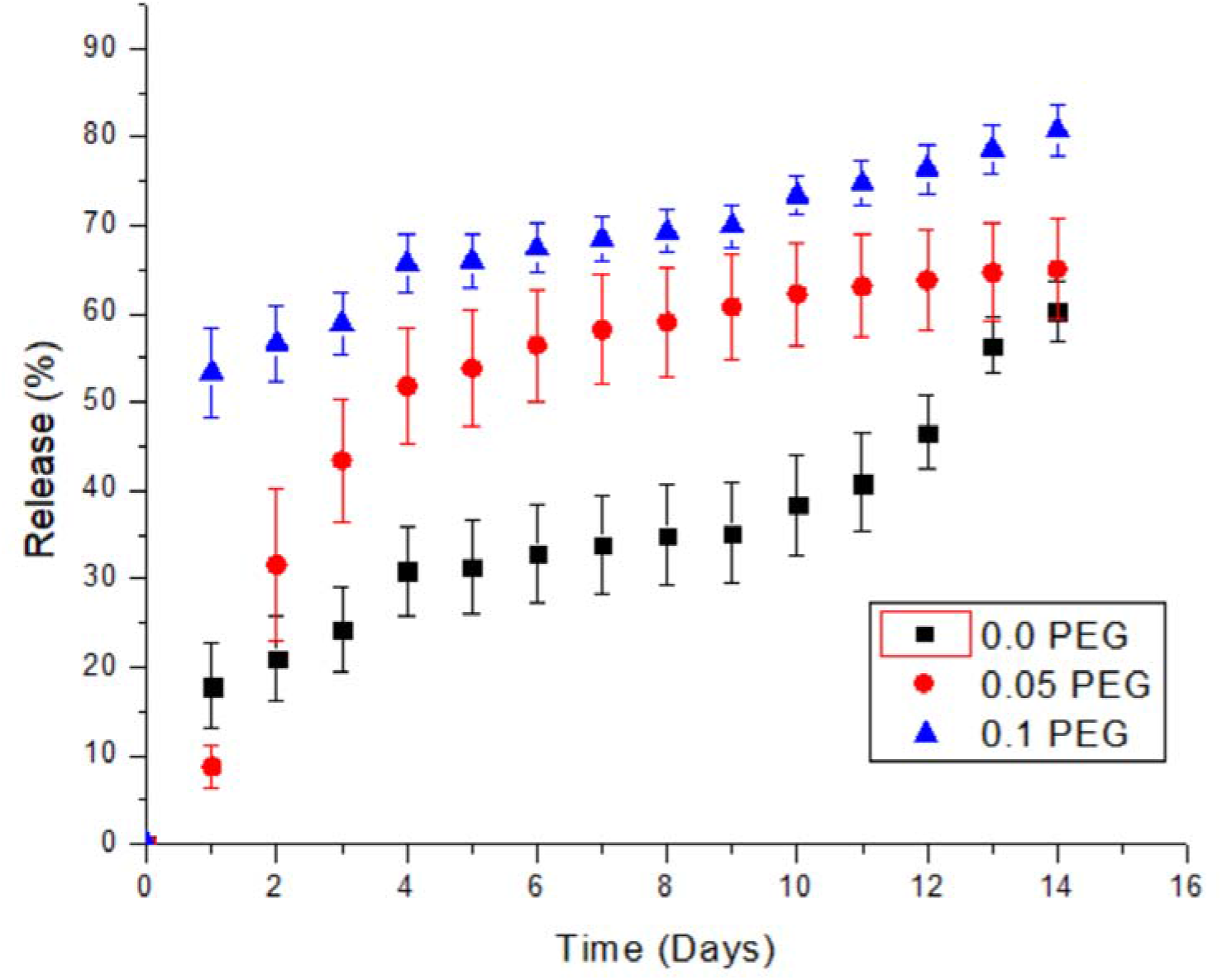
Bev Release in accelerated condition from 0.0, 0.05, and 0.1 PEG PCL implants

**Figure 6** shows a correlation of time frame for the drug release between the accelerated and natural condition. The equation obtained from the graph explains the mathematical relation between time in the two conditions. By multiplying the actual release by a factor of 0.663 (the ratio of accelerated final release and actual final release), the time in PBS and in NaOH can be plotted to find the mathematical relationship between the time in each solution. Using this equation allows for future accelerated condition experiments to be translated to natural conditions without the need to spend 6 months per experiment. The time relation is described in Equation 12.

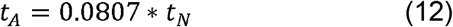

**Figure 6.**
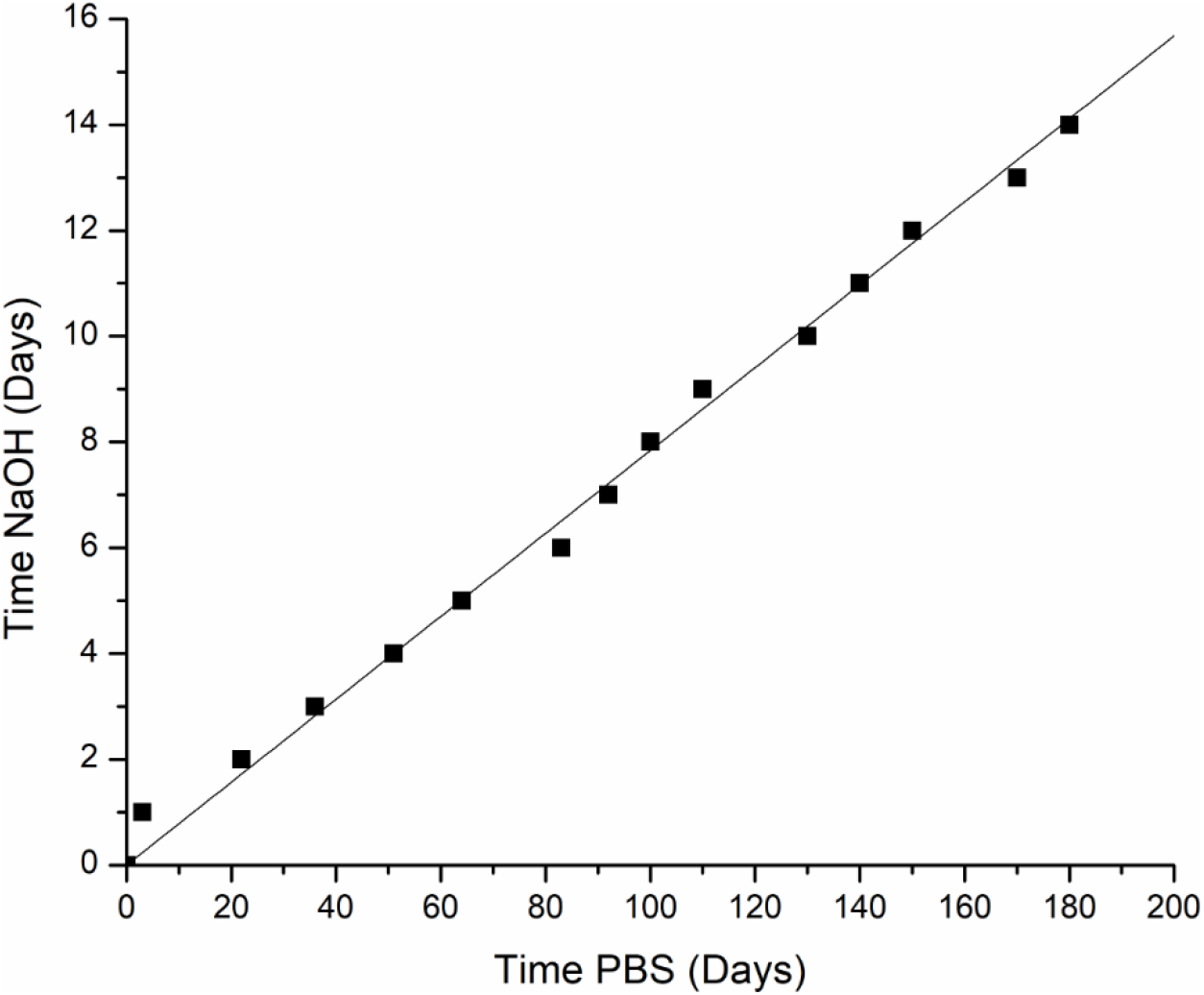
The time correlation between natural and accelerated degradation of the 0.05 PEG PCL capsule.

In this equation, *t_A_* is the time in NaOH and *t_N_* is the time in PBS.

### 3.4. PCL capsule characterization over natural degradation

Based on the SEM images at 2, 4, and 6 months as represented in **Figure 7**, the main effects of degradation are changes in the scale structure over time. As degradation occurred, the scale area dropped in the first month from 5480 ± 990 μm^2^ to 260 ± 50 μm^2^ and then began to increase as degradation continues (p < 0.05) except between month 2 and 3 where there is no significant difference (p > 0.05). The pore size decreased from Day 0 to Month 1 of degradation from 510 ± 30 nm to 370 ± 20 nm (p = 2.35E-5) and then stayed consistent until Month 6 where it increased. The porosity stayed the same for the first month, decreased after 2 months, stayed consistent through Month 4 then increased at Month 5 and stayed consistent through month 6. The trend for 0.05 PEG will be represented as a bar graph later in section 3.5, combined with the accelerated degradation results (Figure 10).

**Figure 7.**
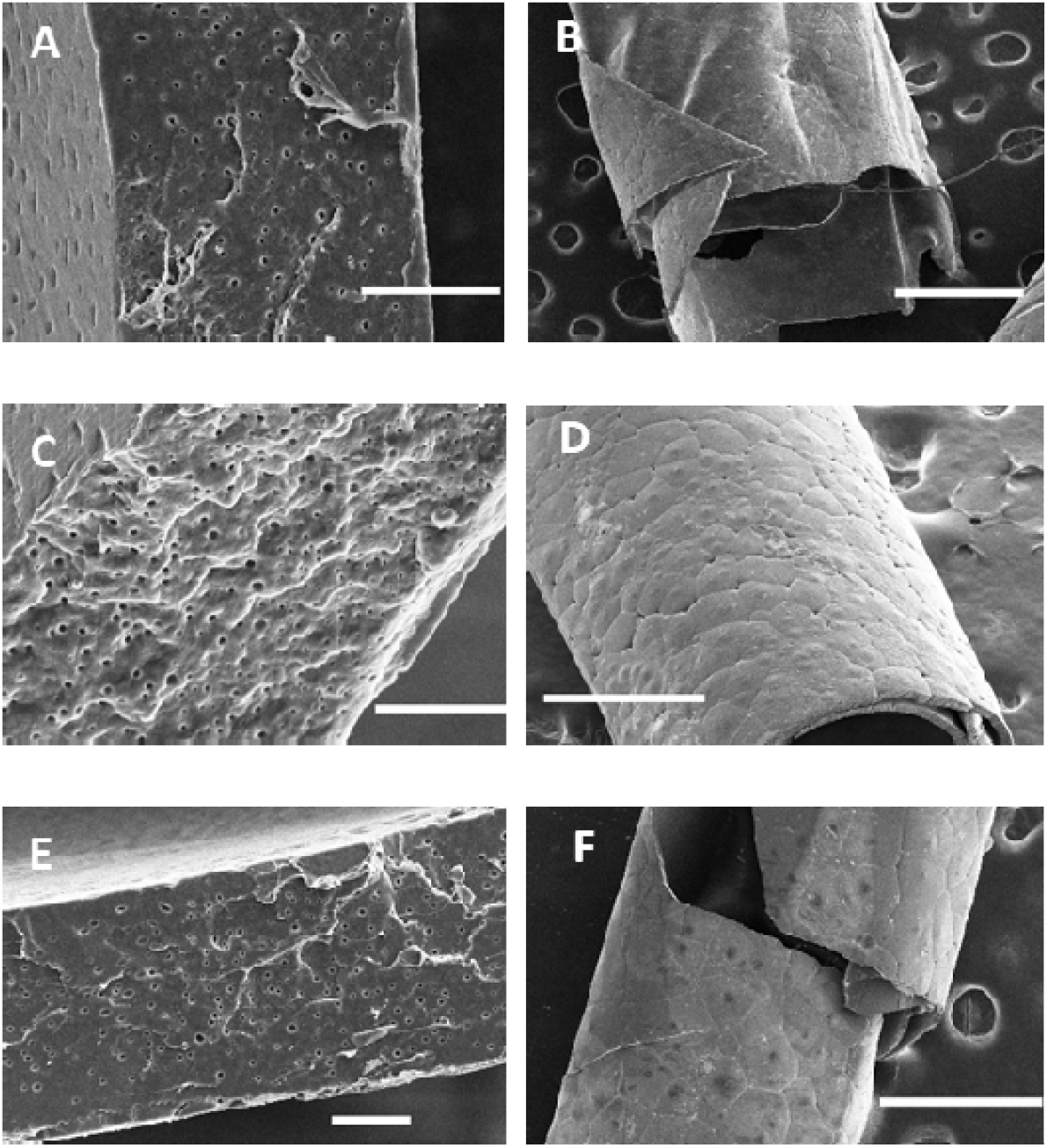
SEM images of 0.05 PEG natural degradation, (A) 2 months cross-section (B) 2 months overall, (C) 4 months cross-section, (D) 4 months overall, (E) 6 months cross-section, (F) 6 months overall. Scale bars = 10 μm for cross-section and 500 μm for overall images.

### 3.5. PCL capsule characterization over accelerated degradation

Using a 0.1M NaOH solution, accelerated degradation was achieved. The 0.05 PEG pore diameter decreased from 510 ± 30 nm to 390 ± 20 nm (p= 1.17E-5) at day 4 of the accelerated condition, exhibited no change by day 10, and then increased at day 20 increasing from 390 nm ± 20 nm to 630 nm ± 20 nm. The porosity did not change until day 10 where it decreased from 3.19 ± 0.097% to 2.22 ± 0.108% (p = 0.011) and stayed steady through day 20. Similar to the natural degradation, the scale area drastically decreases from 5480 ± 990 μm^2^ to 760 ± 90 μm^2^. After 4 days, the scale area increases to 7690 ± 740 μm^2^ before decreasing to 4430 ± 320 μm^2^ at day 20. The thickness decreased in the first 4 days of degradation (p= 3.32E-5), held steady though 10 days, and then continued to decrease through day 20 (p=0.0174).

The 0.0 PEG ratio does not have any pores, thus, there are no measurements for porosity and pore size in Table 3. Measurements for 0.05 PEG 30 days and 0.1 PEG 20 and 30 days have no measurements in Table 3 because significant breakdown of the structure with debris were observed, as seen in **Figure 8F, H and I**. Despite the cracks forming, the 0.0 PEG condition showed no significant changes until 30 days of degradation (**Figure 8G**). After 30 days, the thickness significantly decreased by close to 50% (p=2.73E-4). 0.1 PEG had no significant change in pore size throughout the degradation process, and the only changes were in porosity and thickness. The porosity increased in the first 4 days of degradation from 6.88 ± 0.336% to 9.03 ± 0.121% (p=0.011) then decreased to 5.28 ± 0.267% (p=0.0211). The thickness decreased in the first 4 days (p=0.00129) but did not change significantly between day 4 and day 10 (p=0.211). The trend of scale area and pore diameter is represented in **Figure 10**, integrated with the natural degradation considering the time correlation between the natural and accelerated conditions.

**Table 2.** Pore size, porosity, and thickness analysis from natural degradation SEM cross-section images.

| Month |  | 0.05 PEG |
| --- | --- | --- |
| 1 | Pore Diameter (nm) | $370 \pm 20$ |
| | Porosity (%) | $3.07 \pm 0.089$ |
| | Scale Area ($\mu\text{m}^2$) | $260 \pm 50$ |
| | Sheet thickness ($\mu\text{m}$) | $19.6 \pm 0.23$ |
| 2 | Pore Diameter (nm) | $380 \pm 20$ |
| | Porosity (%) | $1.94 \pm 0.095$ |
| | Scale Area ($\mu\text{m}^2$) | $4130 \pm 970$ |
| | Sheet thickness ($\mu\text{m}$) | $24.2 \pm 0.36$ |
| 3 | Pore Diameter (nm) | $420 \pm 20$ |
| | Porosity (%) | $2.36 \pm 0.059$ |
| | Scale Area ($\mu\text{m}^2$) | $3840 \pm 680$ |
| | Sheet thickness ($\mu\text{m}$) | $22.3 \pm 1.00$ |
| 4 | Pore Diameter (nm) | $340 \pm 20$ |
| | Porosity (%) | $2.08 \pm 0.076$ |
| | Scale Area ($\mu\text{m}^2$) | $8810 \pm 810$ |
| | Sheet thickness ($\mu\text{m}$) | $27.7 \pm 0.38$ |
| 5 | Pore Diameter (nm) | $430 \pm 20$ |
| | Porosity (%) | $3.17 \pm 0.096$ |
| | Scale Area ($\mu\text{m}^2$) | $13870 \pm 2010$ |
| | Sheet thickness ($\mu\text{m}$) | $29.0 \pm 1.22$ |
| 6 | Pore Diameter (nm) | $530 \pm 30$ |
| | Porosity (%) | $4.29 \pm 0.351$ |
| | Scale Area ($\mu\text{m}^2$) | $17790 \pm 2540$ |
| | Sheet thickness ($\mu\text{m}$) | $27.3 \pm 0.85$ |

**Table 3.**
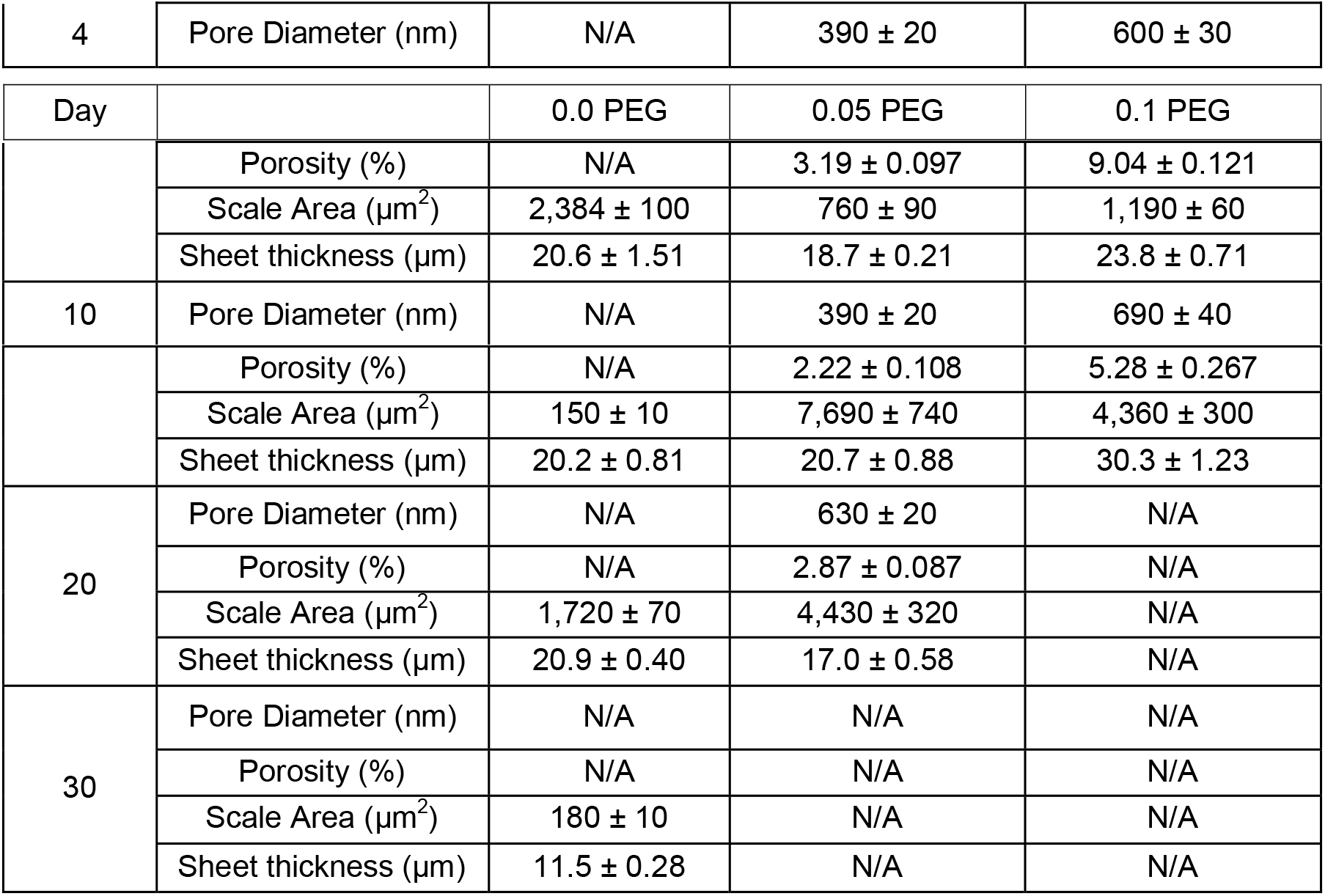
Pore size, porosity, and thickness analysis from Accelerated Degradation SEM cross-section images.

**Table 4.** Values used in Equation 4

| Parameter | Value |
| --- | --- |
| $R_i$ (cm) | 0.0413 |
| $R_o$ (cm) | 0.0464 |
| $L$ (cm) | 1 |
| $K$ | 0.515 |
| $D_e$ ($\text{cm}^2/\text{s}$) | $6.47 \cdot 10^{-11}$ |
| $M_\infty$ (mg) | 1.25 |

**Figure 8.**
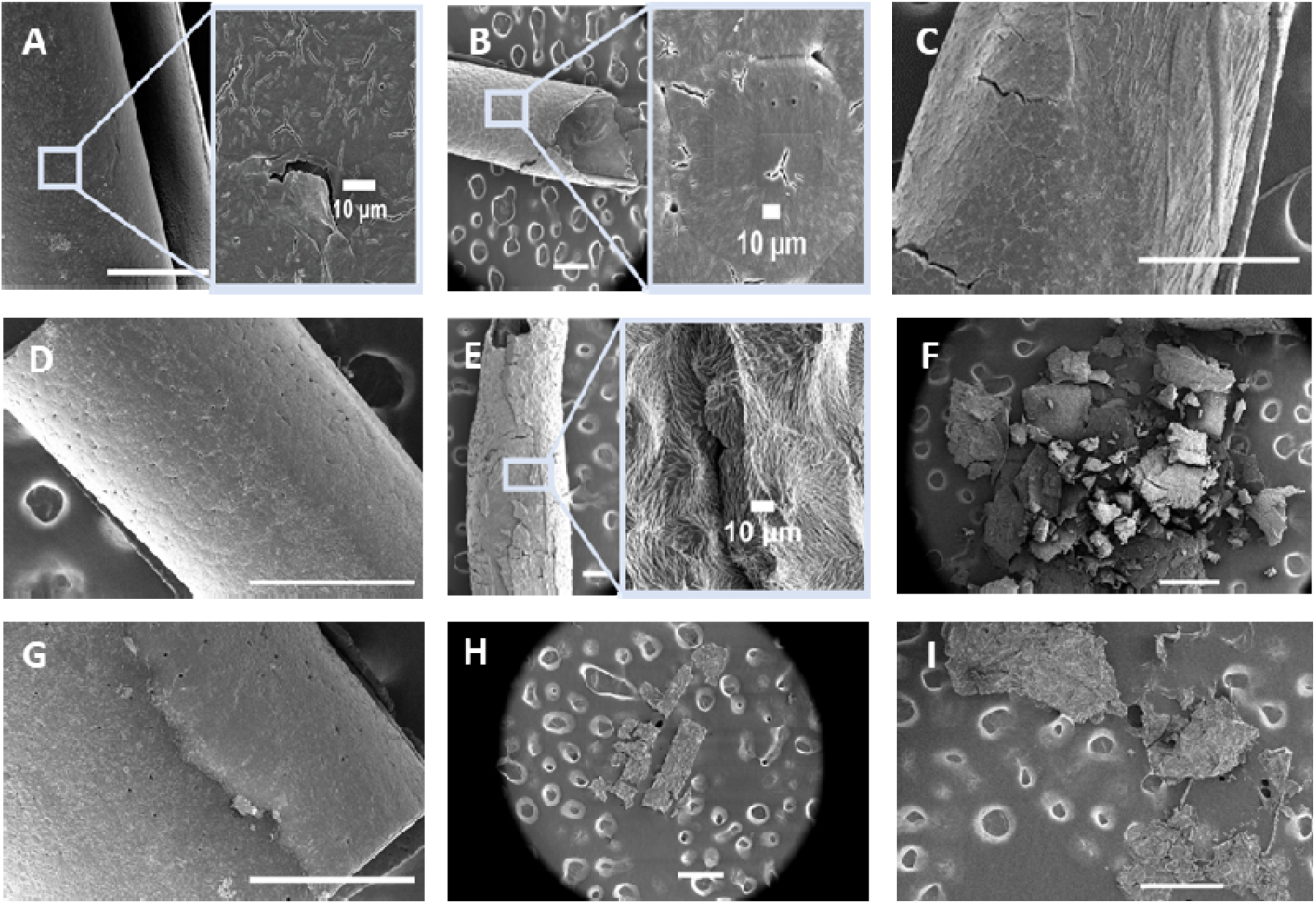
SEM images of Accelerated degradation, (A) 0.0 PEG 4 days, (B) 0.05 PEG 4 days, (C) 0.1 PEG 4 days, (D) 0.0 PEG 10 days, (E) 0.05 PEG 10 days, (F) 0.1 PEG 10 days, (G) 0.0 PEG 20 days, (H) 0.05 PEG 20 days, (I) 0.1 PEG 20 days, (J) 0.0 PEG 30 days, (K) 0.05 PEG 30 days, (L) 0.1 PEG 30 days. Scale bars = 500 μm for all images.

**Figure 9.**
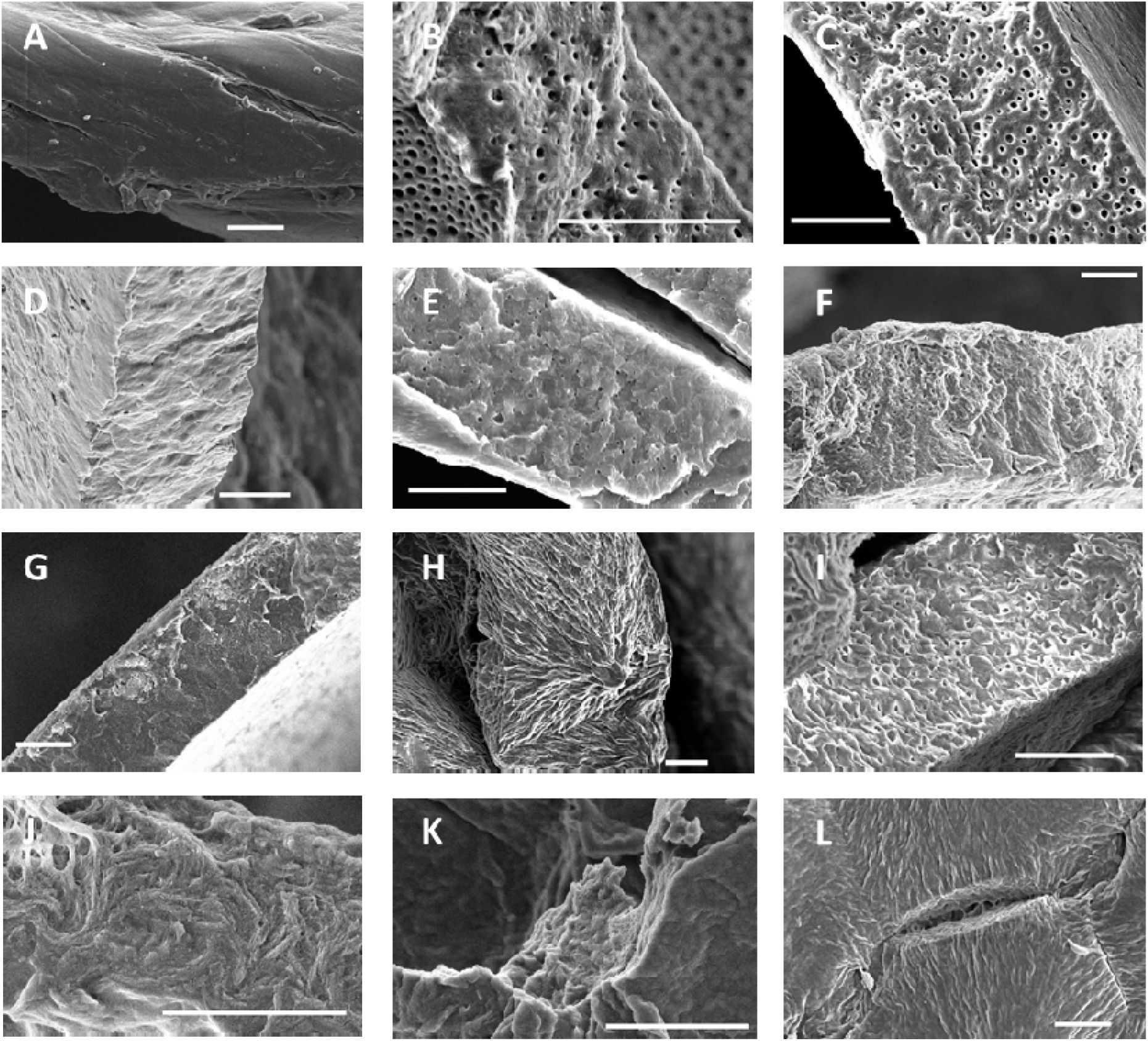
SEM images of Accelerated degradation, (A) No PEG 4 days, (B) 0.05 PEG 4 days, (C) 0.1 PEG 4 days, (D) No PEG 10 days, (E) 0.05 PEG 10 days, (F) 0.1 PEG 10 days, (G) No PEG 20 days, (H) 0.05 PEG 20 days, (I) 0.1 PEG 20 days, (J) No PEG 30 days, (K) 0.05 PEG 30 days, (L) 0.1 PEG 30 days. Scale bars = 10 μm for all images.

Both degradation conditions showed a similar trend in pore size and scale area decreasing after the initial degradation and then increasing at the end to reach the initial pore size before degradation (Figure 10). The scale area at Day 4 in the accelerated degradation condition (red columns) was between the Month 1 and Month 2 scale areas, which lines up with the time analysis (50 days). The scale area at Day 10 was not significantly different from the Month 4 scale area (p = 0.337), which also corresponds to the time analysis of 124 days for 10 days of accelerated degradation. The porosity trend was the same with the initial parts of degradation showing no effect on porosity and then decreasing, but the natural degradation saw an increase at 5 months while the accelerated degradation did not increase before it completely degraded. However, the trend of thickness showed difference between natural and accelerated degradation, probably due to the different degradation mechanisms. The consistent degradation in the natural condition caused no trend to appear in the thickness of the membranes while the increased surface degradation in the accelerated condition caused the thickness to decrease as degradation continued.

**Figure 10.**
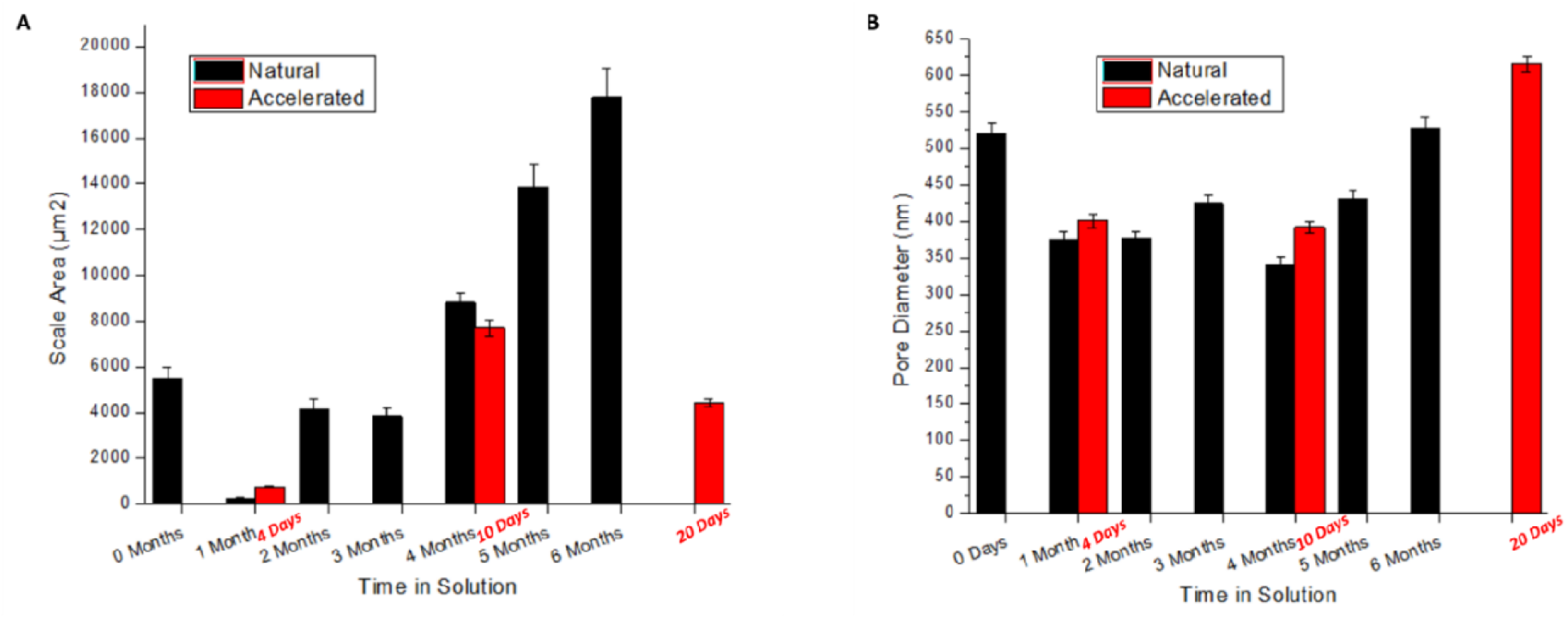
Column graphs of implant characteristics over time in natural and accelerated condition (A) Scale Area (B) Pore Diameter.

### 3.6 Drug release modeling with effective diffusion coefficient (De) and partition coefficient (K)

#### 3.6.1 Effective Diffusion Coefficient (De) and Partition Coefficient (k)

Figure 11 shows the plot of Bev permeating across a 0.05 PEG PCL membrane. For the most part, the permeation is steady state allowing for the use of Equation 2 to determine the permeability coefficient from the slope of the line given by the permeation data. With a donor concentration of 50 mg/mL, an exposed membrane area of 0.78 cm^2^, and 42.5 μg/day permeated across the membrane, the permeability was found to be 1.28E-8 cm/s.

**Figure 11.**
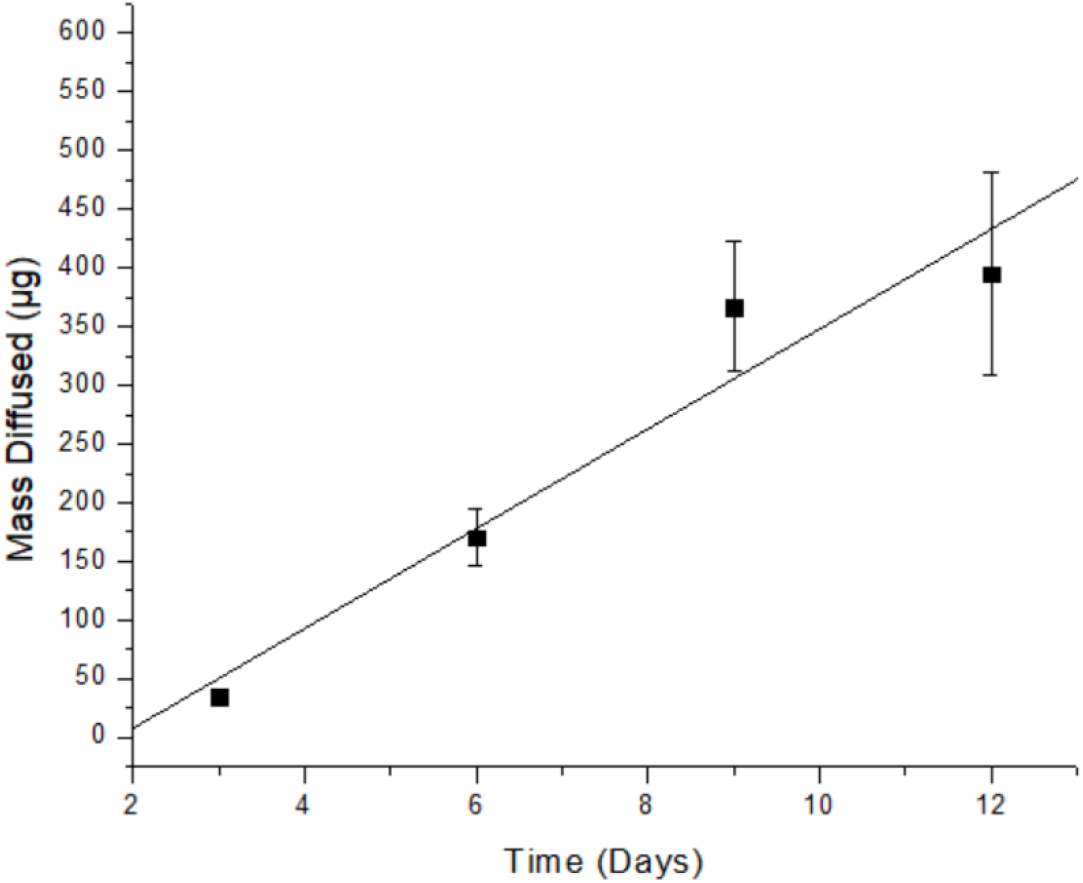
Permeability data: cumulative drug permeation vs. time.

The partition coefficient determined experimentally was K= 0.515 when an equilibrium solution of 20 mg/mL was used. Calculating the Stokes-Einstein diffusion coefficient and hindrance factor gives the final values necessary to use Equation 9 to calculate the effective diffusion coefficient. Using the accelerated degradation images to determine the change in pore size over time, the change in D_e_ over time can be incorporated into the modeling equation for a more accurate representation of the long-term release.

#### 3.6.2 Drug release model

Figure 12 shows the model created from the degradation data and compares it to the actual release data from a 1.25 mg Bev 0.05 PEG ratio implant in PBS. After plotting the model and actual data, the model used fits the actual data almost perfectly. The model uses different De values corresponding to 0, 4, and 10 days of accelerated degradation. Using the time correlation found in Figure 4, at 50 days, the De is from the 4 days degradation, and at 124 days, the 10 days degradation D_e_ was used in Equation 4 to produce the model. As the current model stands, the model predicts 99.56% release by 180 days compared to the 99.71% released in the actual data.

**Figure 12.**
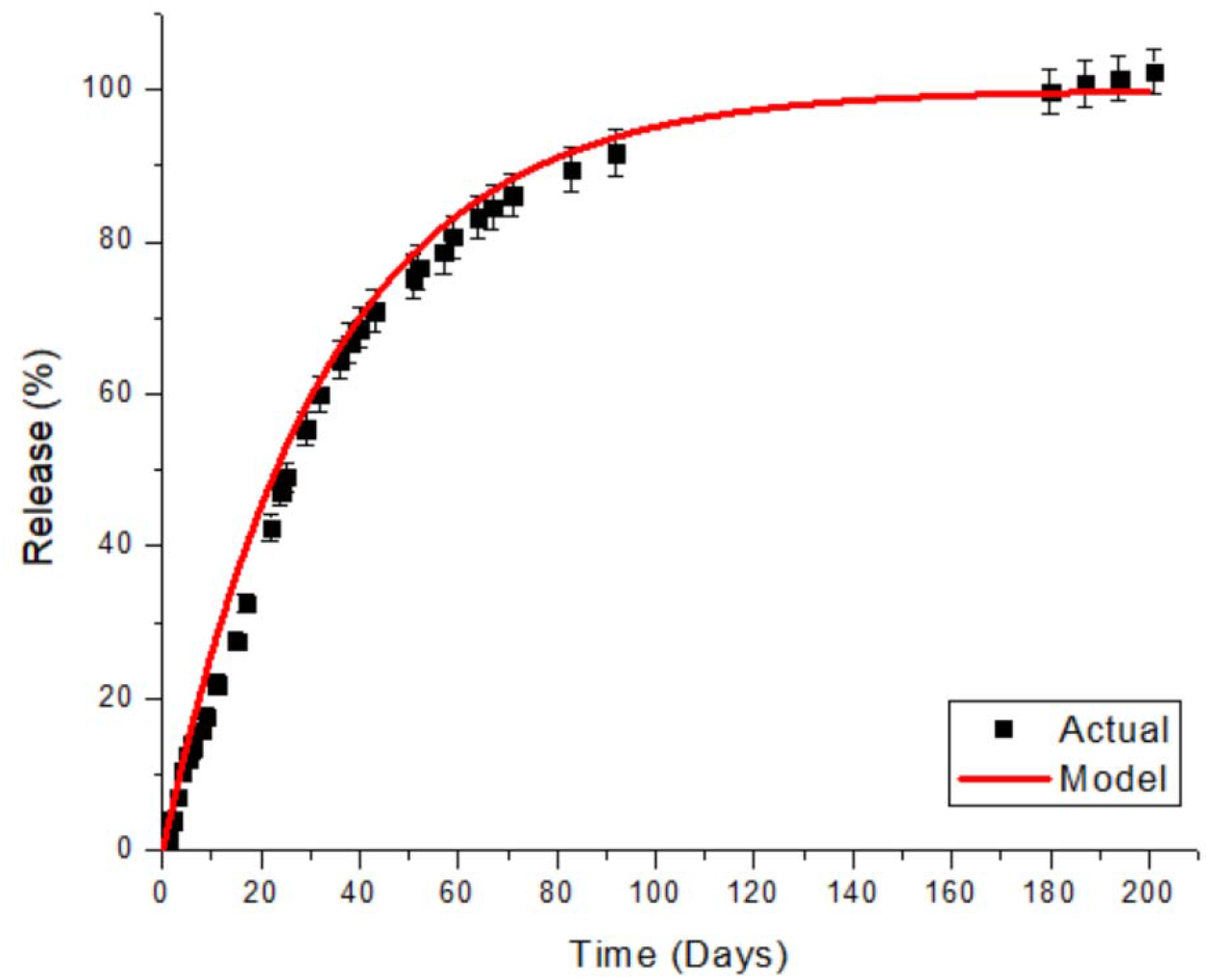
Model fitting for 6 months release in natural condition.

Equation predicted the accelerated drug release kinetics well taking into account the total drug release as shown in Figure 13. Incorporating the time correlation from Figure 4 instead of t and multiplying the result by a ratio of the final release percent for accelerated and natural conditions.

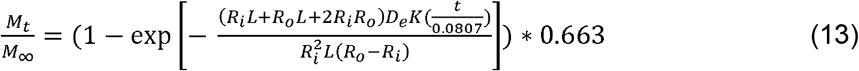

**Figure 13.**
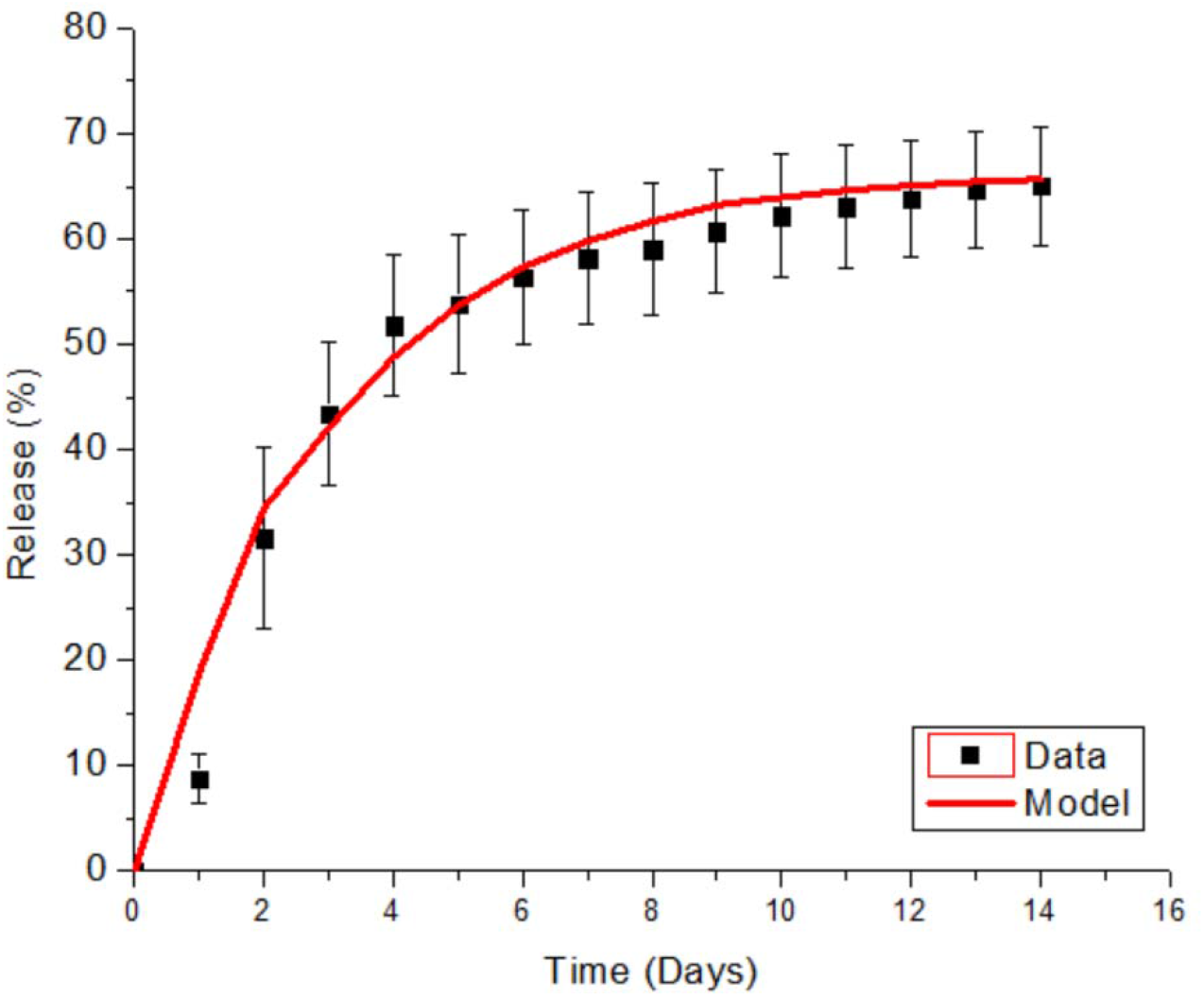
Accelerated Release Model

Equation 13 can be used to model accelerated release to compare how the model in natural condition will represent the actual release in that condition.

## 4. Discussion

After testing the No PEG, 0.05, and 0.1 PEG conditions, the data showed that increasing the PEG concentration in PCL increases the pore size, porosity, thickness of the membrane and the degradation rate. The increased porosity allowed for the natural or accelerated solution to penetrate into the membrane and increase the surface area for degradation and increasing the degradation rate. The decrease in the scale area during the beginning of degradation is most likely due to the breakdown of polymer chains on the surface (**Figure 14**). The initial degradation causes small channels to develop on the larger scales creating a larger number of smaller scales. As degradation continues the area around the small channels eventually levels out creating larger scales again while deepening the remaining ones. The scale structure on PCL sheets was also observed in other publication. ^17^ They mentioned that the scale structure changed over degradation depending on the materials and degree of degradation. The pure PCL’s scale structure disappeared whereas a composite PCL’s structure become more pronounced at 30 days of degradation. This implies that surface degradation or erosion affected the structure.

**Figure 14.**
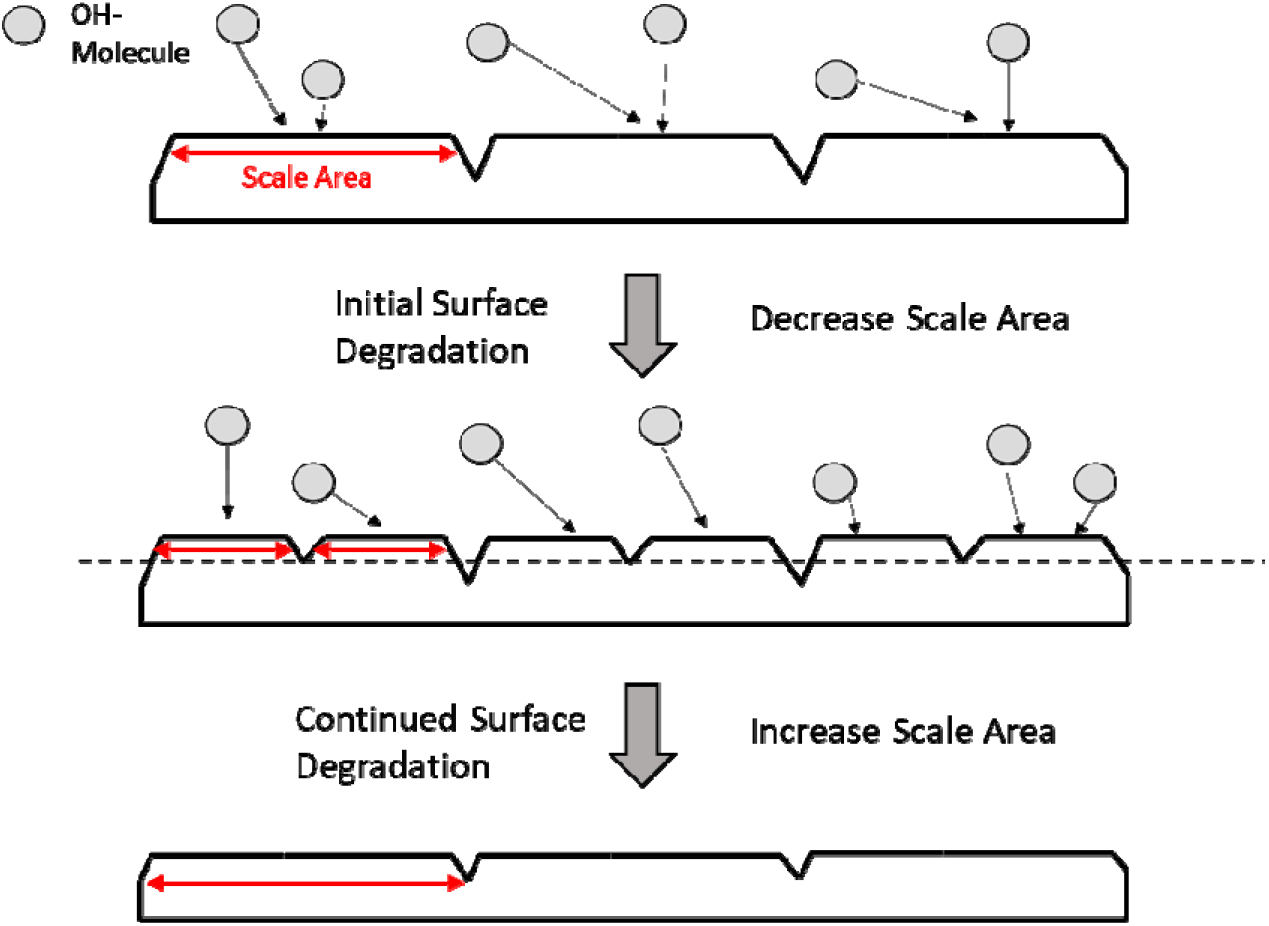
Scale area changes over degradation

The decrease in the scale area during the beginning of degradation is most likely due to the breakdown of polymer chains on the surface (**Figure 14**). The initial degradation causes small channels to develop on the larger scales creating a larger number of smaller scales. As degradation continues the area around the small channels eventually levels out creating larger scales again while deepening the remaining ones. The decrease in pore size can be explained by new smaller pores developing during degradation as well as debris from degradation clogging larger pores. The increase in pore size at the end of degradation can be explained by the merging of the smaller pores to create large ones and the debris being leeched out of the clogged pores. The porosity held steady with the initial degradation because the new smaller pores that formed were offset by the clogging of the larger pores. It eventually decreased due to the clogging and decrease in pore size. The final increase in porosity can be attributed to the PCL between the pores degrading away and debris being leeched out to increase the total volume taken up by the pores.

Correlating the amount of time in PBS and NaOH by the ratio of natural and accelerated final release, a linear relationship between the amount of time spent in PBS and the amount of time in 0.1M NaOH can be determined. This linear relationship was verified by analyzing the change in area for the scale structures present on the implants during degradation. Comparing the 4 day with the 1- and 2-month scale sizes and the 10 day with the 4-month scale size verified the time comparisons made from the linear relationship. Using this relationship, accelerated condition pore diameter and pore size can be used to in the modeling equation to more accurately represent the release kinetics over time without the need of testing the release kinetics for 6 months in natural condition.

The natural condition model accurately predicts the long-term release of Bev from 0.05 PEG PCL implants. With a method of modeling release, any adjustments made to the implant such as adjusting thickness, length, and PEG concentration can be plugged into Equation 4 to see the long-term release. Using initial SEM data, permeability, and the partition coefficient, the long term release can be modeled within a few weeks. Using this model, daily dose can be calculated for varying amounts of loaded Bev and compared to current methods of care to better tailor the release and create effective treatment plans. The accelerated model can be used to compare actual accelerated release and the theoretical release to determine how well the natural model will fit the release data. The accelerated methods used in this study can help expedite the development of biodegradable implants to treat various diseases using mAbs other than Bev.

## 5. Conclusions

We demonstrated monoclonal antibody release for over 6 months using a biodegradable polycaprolactone (PCL) capsule. We identified the drug release kinetics during the period was not affected by degradation because of slow degradation of PCL. However, as observed in the accelerated condition, degradation affecting the breakdown of the overall structure is expected beyond 12 months in physiological conditions. More importantly, we fully understood the longterm drug release kinetics by fitting the first-order kinetics for a cylinder reservoir shape, implying the kinetics can be tuned by model parameters, including dimensions of the capsule, and polymer materials. Lastly, the model can also predict drug release kinetics in the accelerated degradation condition.

## Acknowledgement

This study was partially supported by AMD Award and Lois Hagelberger Huebner Young Investigator Award from Ohio Lions Eye Research Foundation, CCTST, and NIH (R15 EY031500).

